# Landscape structure influences avian species diversity in tropical urban mosaics

**DOI:** 10.1101/388702

**Authors:** Trymore Muderere, Amon Murwira, Paradzayi Tagwireyi, Ngoni Chiweshe

## Abstract

In this study, we tested whether urban landscape structure influences avian species diversity using data for Harare, Zimbabwe. Initially, we quantified landscape structure using fragmentation indices derived from a 5m resolution SPOT 5 imagery. We collected bird species data through field-based observations of birds at 35 locations occurring in five land use/land cover types. We quantified avian species diversity using Barger-Parker, Menhinick and Simpson’s Indices. Regression analysis was used to determine the nature and strength of the relationships between avian species diversity and fragmentation indices. Results indicated that woodland specialist avian species are negatively associated with landscape fragmentation, while grassland specialist and generalist avian species positively responded to patch edge density, habitat patch size and shape complexity. Overall, our results suggest that changes in landscape structure due to expansion of built-up areas in tropical urban areas may influence avian species diversity.

## Introduction

Understanding the factors that influence biodiversity within urban landscapes is fundamental to the planning and development of biodiversity tolerant cities. In the 21^st^ Century, increasing landscape fragmentation resulting from urban development and transportation infrastructure is considered a predominant driver of biodiversity loss in tropical ecosystems [1]. Urban development has a marked impact on the environment [2] as it replaces wildlife habitat with artificial surfaces that are unsuitable as wildlife habitat e.g., asphalt surfaces [3]. Although urban areas occupy <3% of the Earth’s land surface area [4], their ecological impacts span over large spatial extents and sometimes beyond the urban boundaries [5]. Thus, understanding biological diversity-landscape structure (spatial configuration of a given land cover class) relationships is increasingly becoming critical in urban planning [6]. In urban areas, the expansion of built-up areas as well as its configuration is hypothesised to have differential but significant impacts on biodiversity patterns [3], thereby making objective methods for quantifying this phenomena critical.

The quantification of landscape structure in urban landscapes is an important step towards developing urban growth management plans that promote biological diversity. Thus, the development of methods for understanding the impact of urban development on biological diversity in the tropics is critical for biodiversity conservation and enhancement of wildlife persistence in these ecosystems. Such methods may need to focus on improving the estimates of landscape structure-biodiversity relationships. Although field measurements are regarded as the most accurate method of quantifying landscape structure-biodiversity relationships, these measurements are costly and labour intensive and can only be feasible over smaller scales [7, 8]. In this regard the development of methods that supplement field measurements is important.

Developments in Geographic Information Systems (GIS) and satellite remote sensing have made it possible to quantify landscape structure rapidly [2, 3]. In the past, several studies have demonstrated the utility of landscape indices derived from satellite remotely sensed GIS data in estimating landscape-biodiversity relationships across various spatiotemporal scales in temperate landscapes [9-11]. For example, in a study by Coops et al. [12] satellite-derived landscape metrics were used to predict bird species richness in Ontario, Canada using the Moderate-resolution Imaging Spectroradiometer (MODIS) and explained variance ranging between 47 to 75%. Similarly, Guo et al. [10] used a coarse Landsat Thematic Mapper (TM) to estimate avian species habitat relationships in temperate landscapes of Saskatchewan, Canada and their highest coefficient of determination (*R*^2^) was 53%. Wood et al. [11] compared remotely sensed and field-measured vegetation structure in predicting avian species density in Wisconsin, USA and observed that air photo (R^2^ = 0.54) and Landsat TM satellite image (R^2^ = 0.52) were better predictors of avian species density than field-measured vegetation structure (R^2^ = 0.32). In urban landscapes, relatively higher resolution imagery could be of use in modelling the relationship between landscape structure and biodiversity.

The availability of high spatial resolution sensors such as SPOT 5 has provided data that could be used to improve the quantification and mapping of landscape structure indices in urban landscapes that in turn may allow for improved understanding of landscape structure-biodiversity relationships. To date, studies that assess the utility of high spatial resolution multispectral imagery such as SPOT 5 in estimating landscape structure-biodiversity relationships in tropical urban ecosystems remains rudimentary.

In this study, we tested whether and in what way landscape structure indices derived from remotely sensed land cover relate with avian species diversity patterns in Harare, Zimbabwe. Specifically, we tested whether and to what extent avian species diversity respond to constraints including habitat patch size, habitat shape complexity, and habitat inter-patch distance. We derived bird species data from field surveys and landscape structure data from high spatial resolution sensors, i.e. SPOT 5 for Harare, Zimbabwe. We expect differential responses of avian species diversity to habitat constraints. For example, woodland and grassland specialist avian species may be negatively related to decrease in habitat patch size, increased shape complexity and habitat isolation distance. While generalist species will respond positively to changes in habitat conditions.

## Materials and Methods

### Study area

The study was carried out in the Harare Metropolitan province of Zimbabwe (Figure 1). The Harare metropolitan area is approximately 892km^2^ in spatial extent and has a human population of approximately 2.5 million [13]. The center of the study area, is located at Longitude 31^°^7’E and Latitude 17^°^55’S with an altitude range of 1400-1500m above sea level. The city experience two distinct seasons i.e., hot wet summers (October – April) and cool dry winters (May – September). The mean annual rainfall ranges between 800-1000mm, while mean annual temperature ranges between 25 – 27 ^°^C [14].

**Figure 1.**
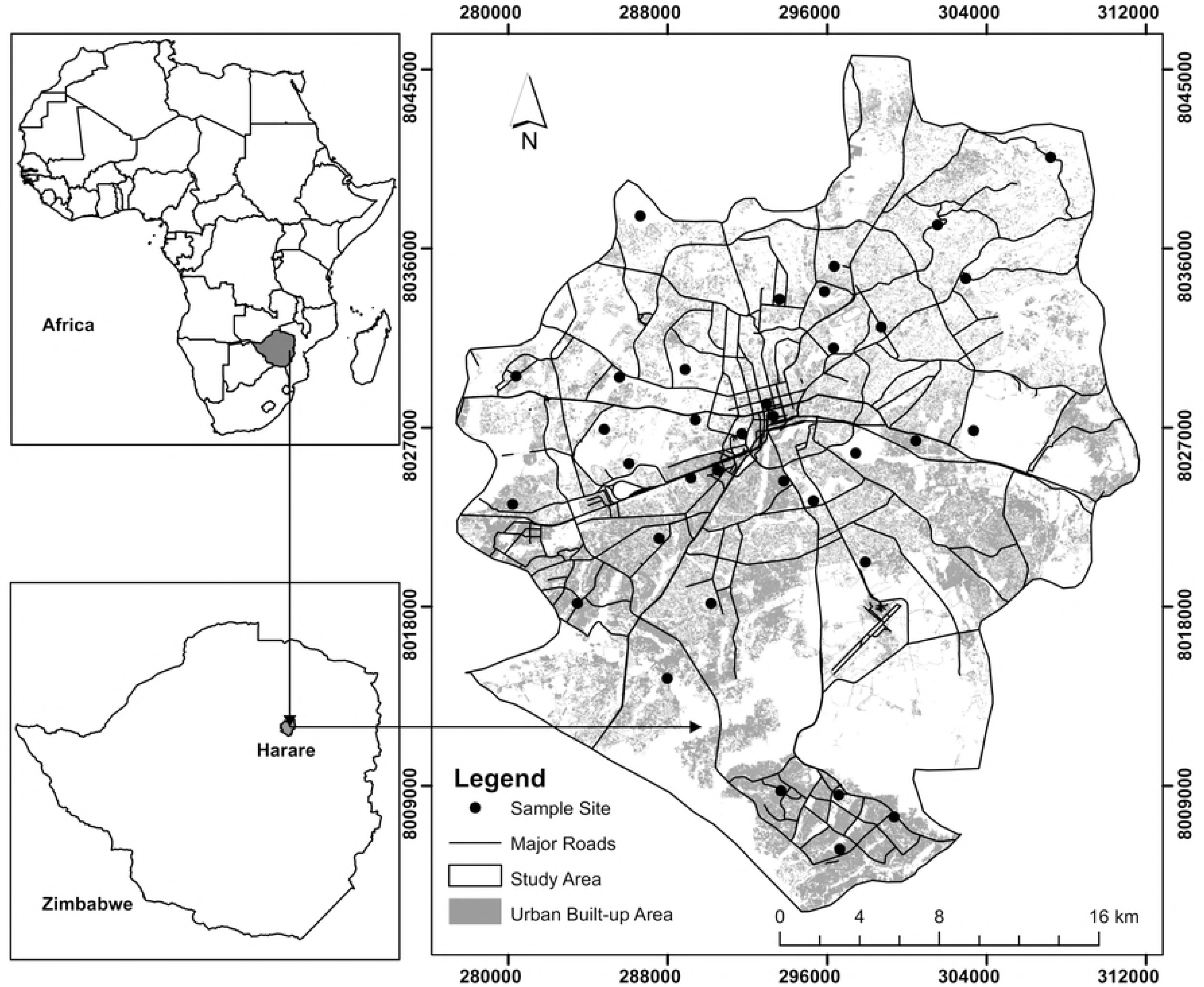
Map of the study area showing the 35 bird observation sites georeferenced in WGS84 and the coordinates are in Decimal Degrees

Our own fieldwork showed that the prevalent land use/land cover (LULC) types in the city include grasslands/pasture and cropland (64.0%), forested (21.0%), urban built-up areas (10.7%), bare ground (3.8%) and water (0.5%). The forested land cover type is mainly deciduous dry Miombo woodland dominated by *Brachystegia spiciformis, Julbernardia globiflora* and *Uapaca kirkiana* [15]. The bare ground cover type consists of exposed surfaces and area under active urban development. The water cover type includes impoundments and rivers. The urban built-up area is made up of impervious surface covering including road networks, industrial areas, high and low density residential areas. The study site was selected because it represents an ideal location to study landscape structure-biodiversity relationships in the context of regional and urban planning. The area is currently undergoing a rapid increase in human population associated with unguided urban development patterns whose impacts have not been quantified.

### Quantifying landscape structure

We derived landscape structure data from a 5-m spatial resolution SPOT 5 image of Harare. Specifically, using Object Based Image Analysis (OBIA) in Trimble eCognition (Trimble, Munich, German) on a desktop computer, we obtained discrete landscape classes of habitat patches for avian i.e., (1) forested areas, (2) grasslands as well as (3) built up areas. Overall mapping accuracy was 89.7%, Kappa coefficient of 84.3% based on 340 sampling test points. We used the Effective Mesh Size [16, 17], grid mesh (mesh size = 4000m^2^) to characterize the landscape structure in the study area. The 4000m^2^ mesh size was used because this represents the average home range size of typical urban birds [9, 16, 18-21]. We then used the Patch Analyst tool [22] in ArcGIS 10.2 (Environmental Systems Research Institute, Redlands, California, USA) following Tagwireyi and Sullivan [23] to quantify landscape structure (configuration and composition) based on 16 landscape patch metrics default in the Patch Analyst tool and 2 Effective Mesh Size landscape patch metrics default in the Effective Mesh Size tool [16, 17]. We tested the 18 patch metrics for multi-collinearity with pairwise Pearson’s correlation [24] and removed all metrics with *R^2^* > 0.90 from further analysis following Graham [25]. Patch metrics were highly variable across landscape classes (SI 1).

### Sampling design

In a GIS, we processed the study area into a LULC categories layer representing three LULC types i.e., low urbanization grasslands, low urbanization forested area and built-up areas (Table 1). Subcategories were defined for each category to account for variations each context presented. Altogether we had seven LULC subcategories and representing three LULC types and 35 transect sampling sites (Table 1).

**Table 1.** Study sites sampling design matrix and definitions of land cover/land use types

| Land Cover/Use Type | Subcategory | Description | Transects |
| --- | --- | --- | --- |
| Low urbanization grasslands | 1 | Grasslands in Low Density Residential Areas | 5 |
|  | 2 | Grasslands in High Density Residential Areas | 5 |
| Low urbanization forested areas | 3 | Forested areas in Low Density Residential Areas | 5 |
|  | 4 | Forested areas in High Density Residential Areas | 5 |
| Built-up area: | 5 | Low Density Residential Areas | 5 |
|  | 6 | High Density Residential Areas | 5 |
|  | 7 | Central Business Districts and Industrial Areas | 5 |

Using the LULC categories base map and the Random Sampling Tool in Quantum GIS 2.6.1 (QGIS Development Team, Switzerland) we stratified the study area (excluding private and security areas e.g., military and airport land) into five sampling sites for each LULC subcategory (total 35 sites) (Table 1). We deemed the sample of 35 sites representative for statistical purposes following Rawlings et al. [24]. Each of the points was used as the center of the 600m transect lines along which we surveyed the birds. The sampling sites were positioned at least 1.5 kilometers apart to ensure spatial independence between surveyed avian species and on different land cover types to account for habitat variation within sites [26].

### Avian species surveys

At each sampling site we recorded observations of diurnal-active birds using an effective detection distance of 50m [27] along either side of the 600m sampling lines. The surveys were done at four different times of the day i.e. between: 6am-9am; 9am-12pm; 12pm-3pm; 3pm-6pm during the summer months of February and April 2015 (hot-wet season), to account for differences in avian species behavior on different times of the day [20, 28]. On each visit, the same observers waited for about five minutes to allow avian species to resume normal activity following MacArthur and MacArthur [28] and then recorded all avian species seen patched, flying or foraging within a 50m distance from the 600m transect line (see SI 1). We identified the birds to species level based on expert knowledge and a field guide book i.e., Roberts Birds of Southern Africa [29]. We also categorized avian species into three ecological guilds (generalists, woodland specialists as well as grassland specialists) because we investigated landscape influence on the birds at guild level.

Avian species were selected as the model species, because they are highly mobile and can respond to landscape change quickly than ground dwelling mammals or other rarely seen species [9] which makes birds useful indicators of species responses to urban development induced environmental change. The study focused on overall avian species than select target species, common in many studies [30]. The advantage of focusing on overall avian species is that it allows the study to account for avian species with different life histories and behaviors [18, 30, 31].

### Quantifying avian species diversity

We used the Menhinick, Berger-Parker and Simpson’s indices to quantify avian species diversity [9, 32, 33] (Table 2). A diversity index is a mathematical measure of biodiversity providing important information about rarity and commonness of a species in a community [33]. We calculated Menhinick’s Index as 1-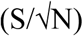 where N= the number of individuals in a sample and S = the number of species recorded [34]. The Berger-Parker Index was estimated as 1-N_max_/N, where N_max_ is the number of individuals in the most abundant species and N is the total number of individuals in a sample [32]. The Simpson’s Index was estimated as 1-∑P^2^, where P^2^ is the total number of organisms of each particular species from the total number of organisms of all species [33]. We chose the Menhinick, Berger-Parker and Simpson’s Indices because they are spatially and temporarily stable, robust and biologically intuitive measures of biodiversity [34], although they remain susceptible to sampling size [33, 35]. We applied the reciprocal 1-D to the indices so that an increase in the index accompanies an increase in diversity for ease of intuitive interpretation following Whittaker [36] and Magurran [34].

**Table 2.** Summary statistics of bird species observed by landscape type

| Land use/cover class | Abundance | Richness | Menhinick's Index | Berger-Parker Index | Simpson's Index |
| --- | --- | --- | --- | --- | --- |
| Low urbanization forested | 1286.00 | 25.00±36.00 | 2.45±2.88 | 0.78±0.86 | 0.90±0.93 |
| Low urbanization grasslands | 1980.00 | 28.87±11.21 | 1.64±0.34 | 0.14±0.82 | 0.80±0.90 |
| Built-up Area | 2815.00 | 18.13±11.68 | 1.60±2.29 | 0.70±0.15 | 0.95±0.06 |

### Relating landscape fragmentation indices to avian species diversity

Prior to regression analysis we tested the avian species data for normality using the Kolmogorov-Smirnov test to test [37] for conformity to the simple regression assumption for randomness and we found a normal distribution (*p*>0.05). We then used simple regression analysis to examine the direction and strength of the relationship between fragmentation indices (independent variables) and avian species diversity (dependent variables) in MS Excel and Statistical Package for Social Science Version 18 [38]. The strength of each regression model was evaluated based on the coefficient of determination (*R*^2^) and the level of significance (*p*-value).

## Results

### Avian species diversity-landscape structure relationships

We surveyed 6081 birds representing 69 species in 35, 600m transects. Thirty percent of the surveyed birds were observed in low urbanization grassland habitat, 46% in built-up areas and 24% in low urbanization forested land. We also observed that bird species abundance, richness and diversity (i.e., Menhinick’s, Berger-Parker and Simpson’s Indices) varies across the three LULC classes (Table 2).

### Woodland specialist avian species - landscape structure relationships

Simple regression showed that woodland specialist avian species were negativity associated with patch metrics derived from low urbanization forested cover type, specifically shape complexity (*R^2^* = 0.635), shape size (*R^2^* = 0.616) and isolation distance (*R^2^* = 0.778) (Figure 2).

**Figure 2.**
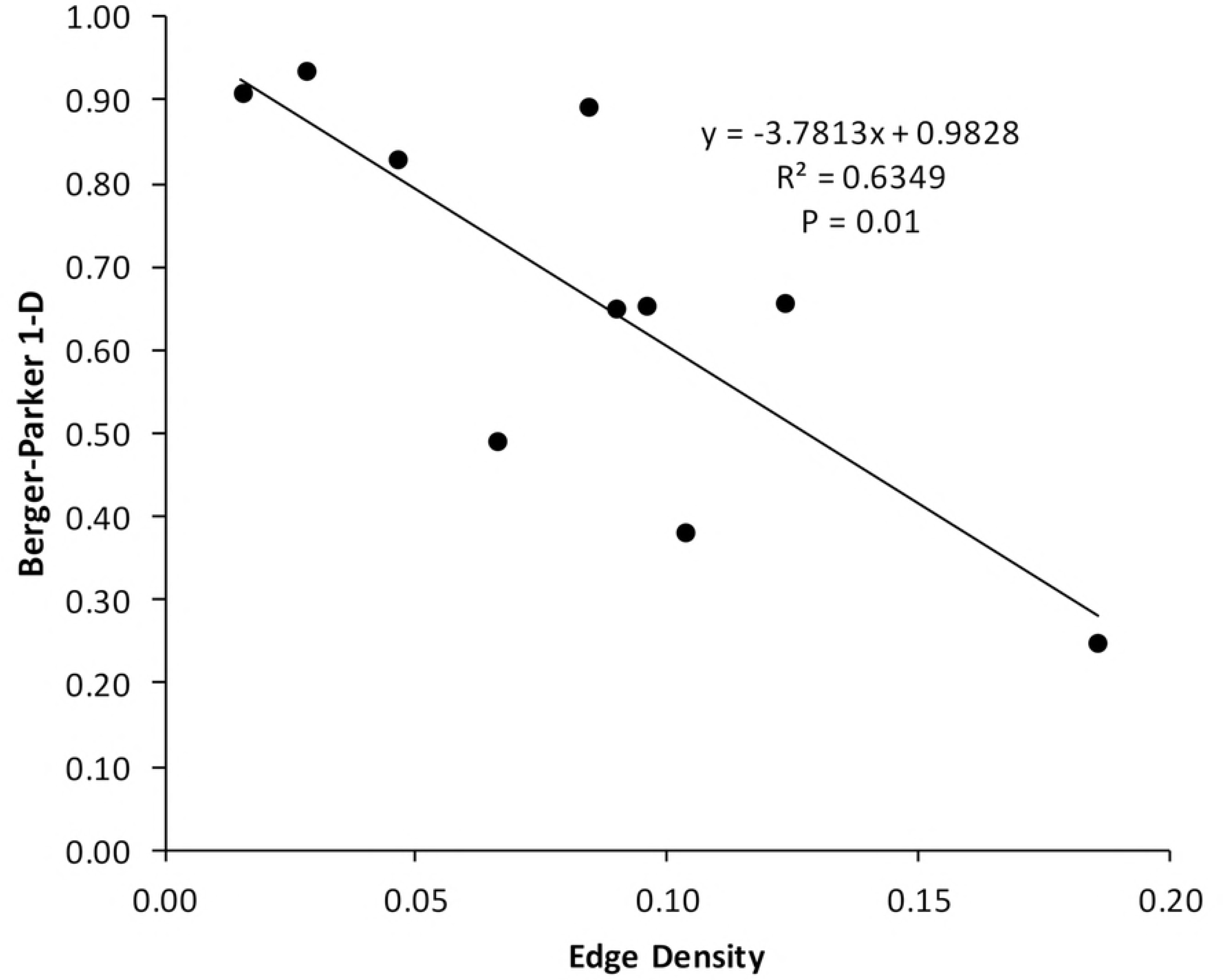
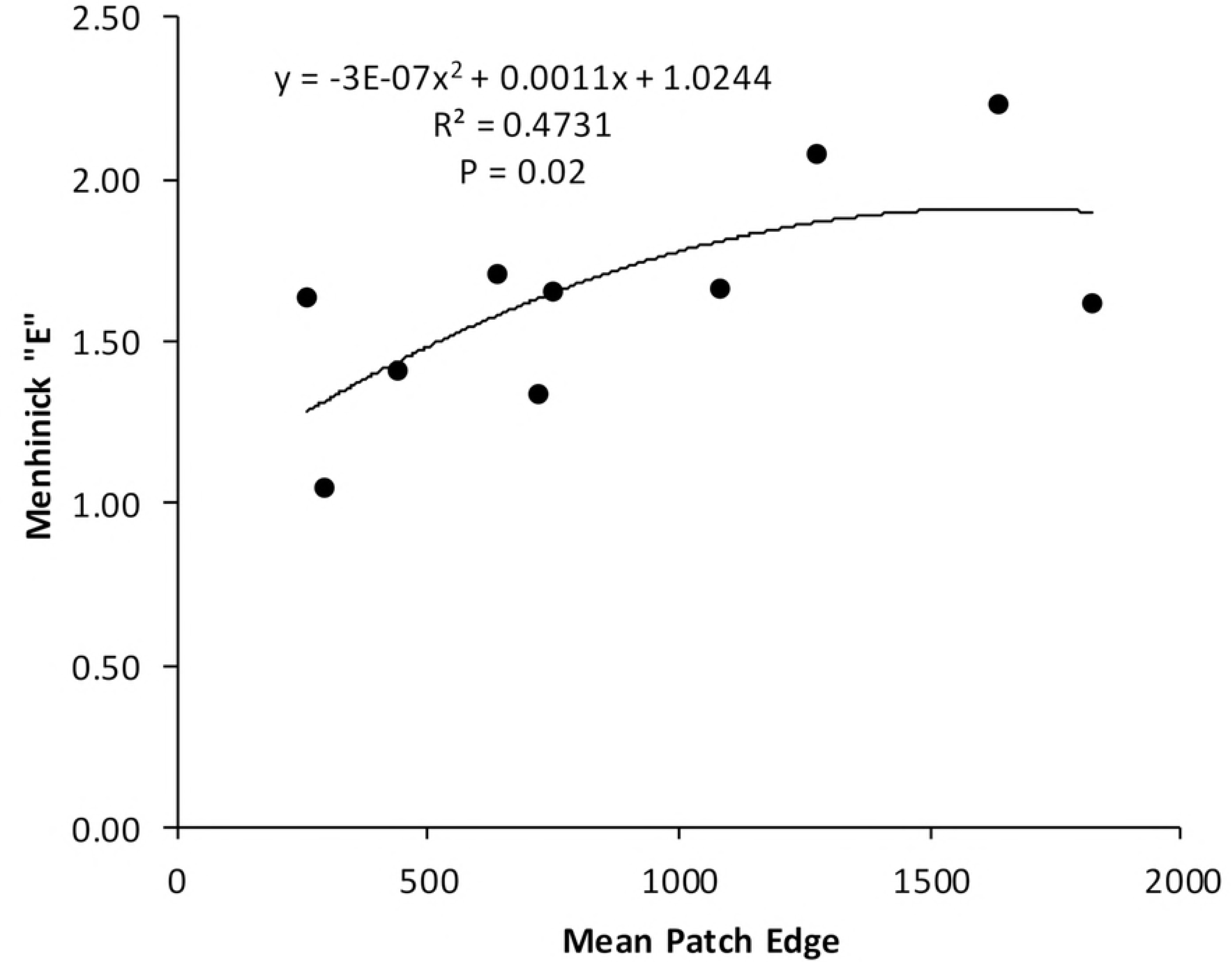
Relationship between woodland specialist avian species diversity (Menhinick’s, Berger-Parker’s Indexes) and landscape structure (a = size, b= shape, c = isolation distance)

### Grassland specialist avian species - landscape structure relationships

Simple regression showed that grassland specialist avian species had a strong positive polynomial relationship with patch edge derived from low urbanization grassland cover type (*R^2^* = 473, Figure 3) and not significant (*p>0.05*) association with patch size and isolation distance.

**Figure 3.**
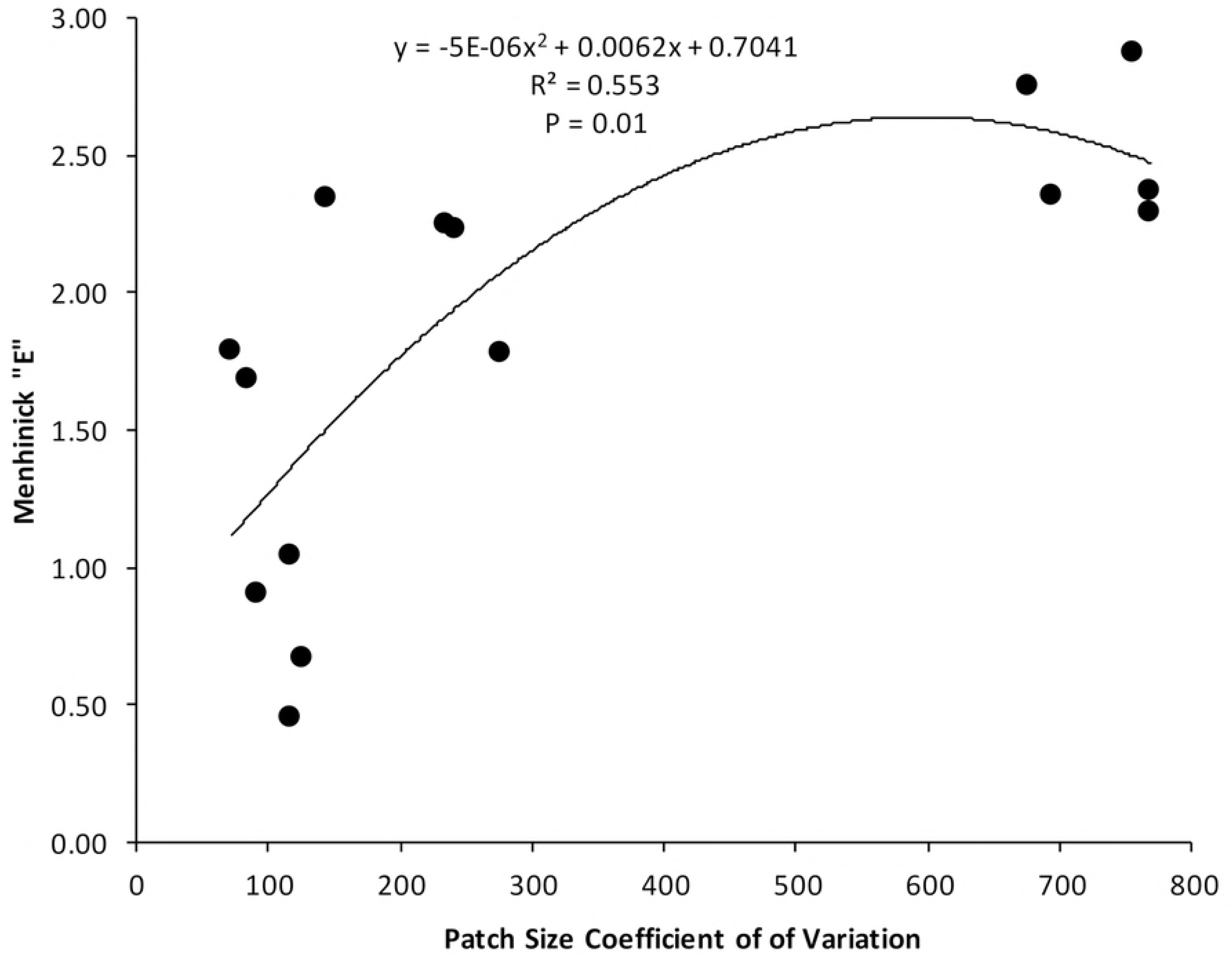
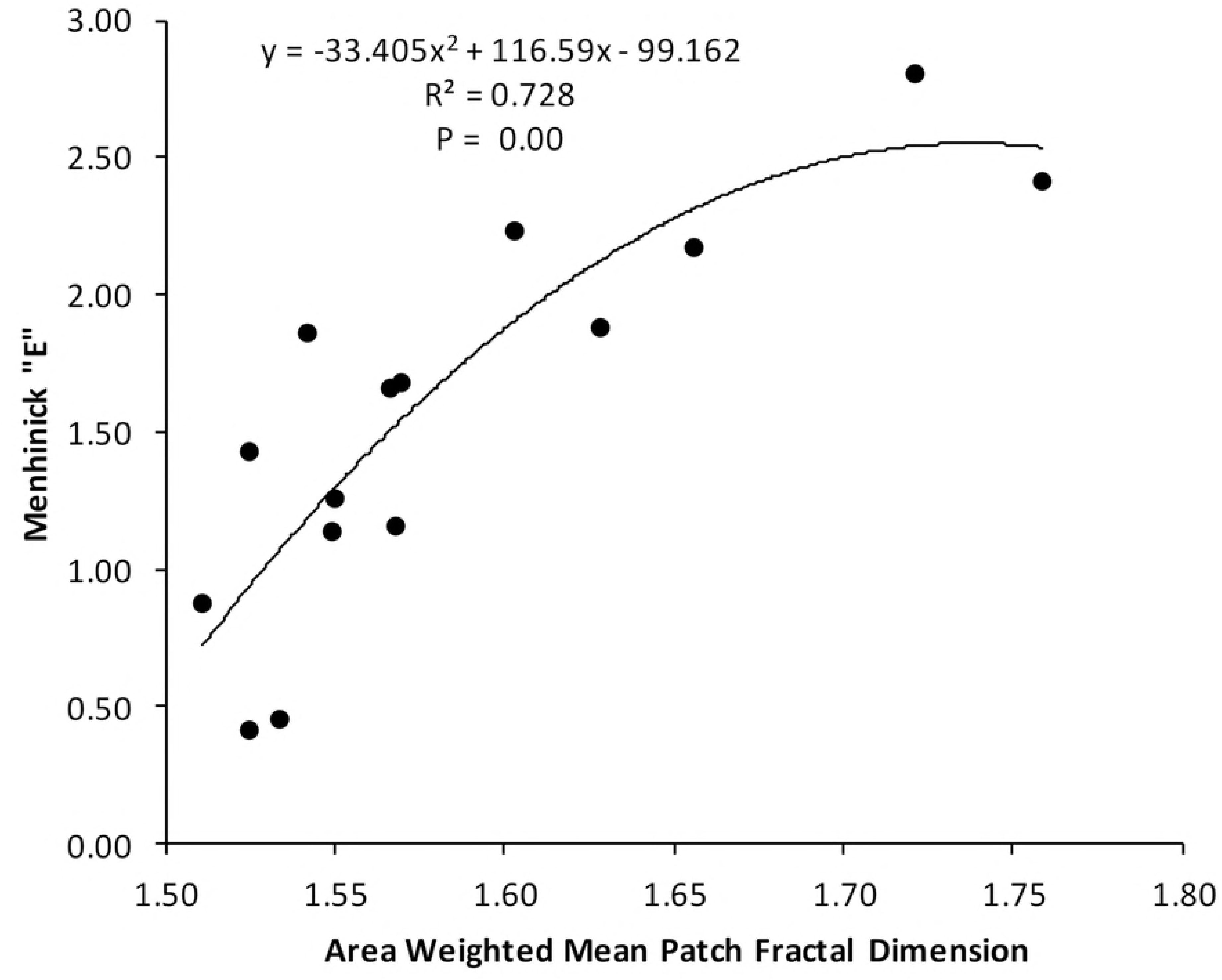
Relationship between grassland specialist avian species diversity and landscape structure (shape)

### Generalist avian species - landscape structure relationships

Simple regression showed significant (p <0.05) positive regression between generalist avian species and habitat fragment size, shape complexity (*R^2^* = 0.553, *R^2^* = 0.728) (Figure 4) but not significant relationship with isolation distance of the intensely built-up cover type.

**Figure 4.**
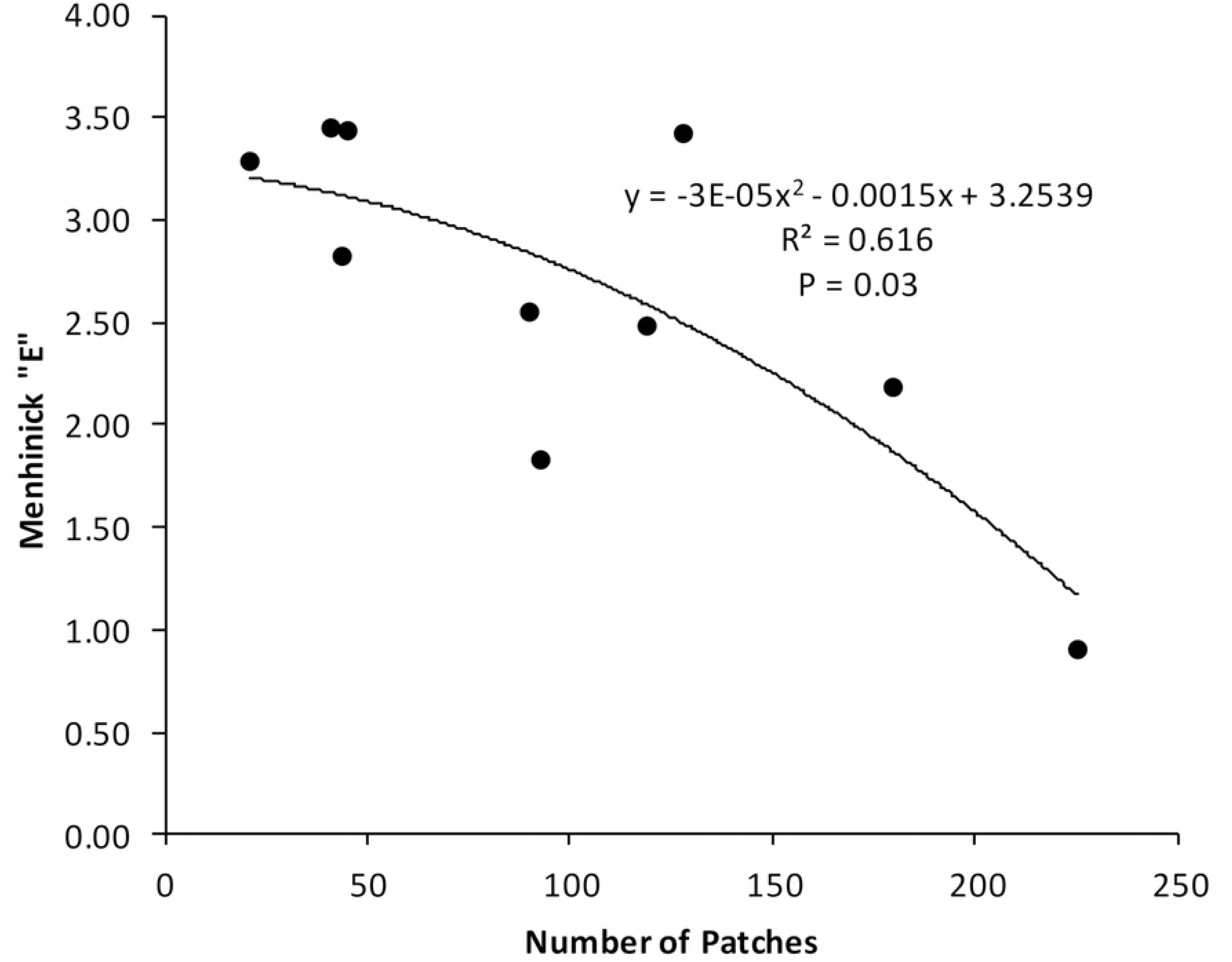
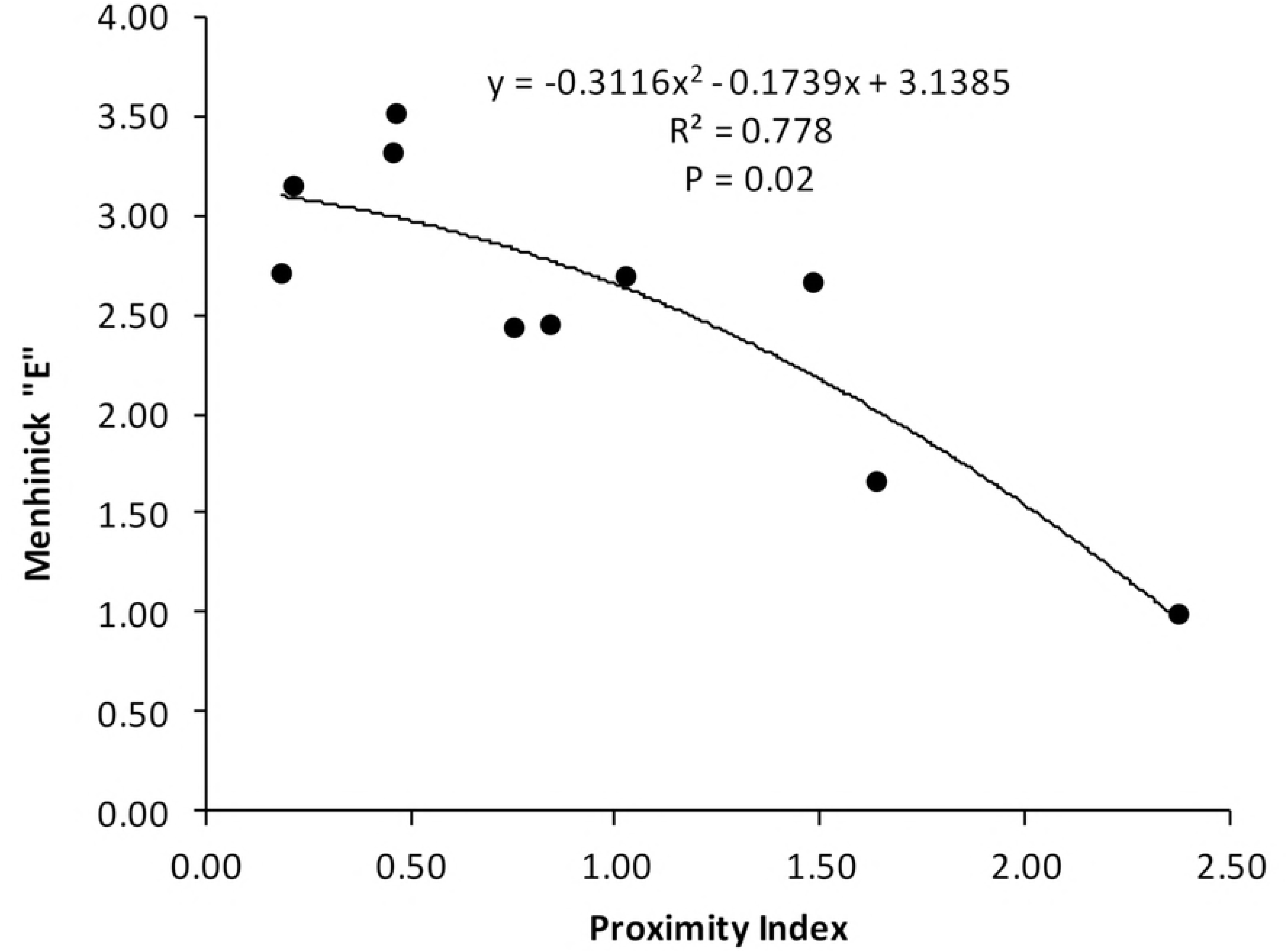
Relationship between generalist avian species diversity and landscape structure (a= size, b= shape)

## Discussion

Results of this study indicate that landscape structure elements influence avian species diversity in the study area. These results are consistent with our initial hypothesis that landscape structure influences avian species diversity in urban landscapes. The results are also consistent with findings of previous studies in urban and non-urban landscapes of North America [e.g.,39], Central Europe [e.g.,18] and Australia [e.g., 40] who observed that landscape constraints operating at habitat level influence avian species diversity.

Results also indicated that avian species diversity of woodland specialists negatively correlated with edge density of the low urbanization forested cover type, suggesting that for these specialist avian species increased fragmentation in woodlands due to urban development has negative impacts on them. This is not surprising as McWilliam and Brown [39] also observed similar responses in Ontario, Canada where a decrease in forest cover size accompanied by an increase in the size of built-up area caused decline in the diversity of forest interior specialist species over a ten-year period. Rodwald and Yahner [41] also observed linkages between landscape composition and avian community structure in central Pennsylvania, USA. However, the result is significant in informing urban planning practices that may need to preserve woodland specialist species. We therefore deduce that landscape metrics derived from high resolution imagery can be used for accurate estimation of avian species diversity in urban landscapes. In contrast, but not surprising, results also indicated that grassland specialist avian species positively correlated well with habitat shape complexity especially high edge effects. These results suggest that while woodland specialists are negatively affected by woodland fragmentation, this process facilitates grassland avian species expansion. Again this result is consistent with Jones and Bock [42] who reported that open spaces typically low urbanization grassland areas can sustain a high diversity of grassland avian species. Although this is not surprising this result is important for aiding urban planning practices that may need to conserve various bird species with different habitat preferences.

The observation that generalist avian species diversity positively correlates with landscape fragmentation also suggest that generalist bird species benefit from forest loss and fragmentation. This is consistent with previous studies from Central Europe [e.g., 21, 35, 43] and North America [e.g., 6, 18, 31, 44] which link the behavioral traits of generalist avian species to ubiquitous opportunities presented by intensely built-up landscapes.

Overall, this study provides evidence that high resolution satellite imagery offer improved opportunities for estimating the effect of urban development on biodiversity in particular avian species diversity. The best model explained 79% variation in avian species diversity. This coefficient of determination is higher than obtained by Coops et al. [12] and Guo et al. [10] across various spatiotemporal scales in temperate landscapes. Coops et al. [12] used a number of vegetation indices derived from MODIS to predict breeding bird species richness in Ontario, Canada and their highest coefficient of determination (R^2^) was 75%. Guo et al. [10] and Wood et al. [11] on the other hand found weak to average relationships between landscape spectral vegetation indices and avian species diversity derived from a coarse Air photo and Landsat Thematic Mapper (TM) to estimate avian species habitat relationships in temperate landscapes of Saskatchewan, Canada and Wisconsin, USA respectively and their highest coefficient of determination (*R*^2^) was 54%.

This study differs from previous studies in three main ways. Firstly, studies that have used vegetation to estimate avian species diversity in the temperate regions have used medium to low spatial resolution imagery data such as Aerial, Landsat and MODIS images. These factors have resulted in weak relationships, high errors and uncertainties. However, our study estimated avian species diversity from landscape metrics derived from high spatial resolution satellite imagery with very low error margins. Thus, it is important to note that integrating landscape metrics derived from high spatial resolution satellite imagery improved avian species prediction compared to previous studies. Secondly, there is paucity in studies conducted in tropical ecosystems that relate landscape metrics to avian species diversity in urban landscapes yet avian species diversity is a biodiversity indicator that has important insights to the science of urban environmental change. Finally, unlike previous studies that only determined the relationships between vegetation indices and avian species diversity we quantified landscape structure attributes in terms of size, shape and isolation distance at a fine spatial scale. Again, we find this especially important in African tropical urban landscapes where tree cover is low, much of the built-up areas have no tarmac cover and much of the urban development is informal and poorly planned, thus making high spatial resolution satellite imagery an excellent alternative to delineating spatial variability habitat fragmentation. However, it will be useful to test the applicability of these models in independent study sites to observe whether the form of remotely sensed models of landscape metrics are consistent and can be improved further. Nevertheless, we make a claim that this finding provides an opportunity to quantifying the impact of urban landscape pattern on biodiversity in tropical urban landscapes of sub-Saharan Africa.

## Conclusion

The main objective of this study was to test whether and to what extent avian species respond to constraints including habitat fragment size, shape complexity and isolation distance in urbanizing tropical ecosystems. From the results of this study, we conclude that the:

1. size, shape and isolation distance of habitat fragments matter to woodland specialist avian species;
2. shape of habitat fragments matter to grassland specialist species, than isolation and size of grassland fragments; and
3. the increasing complexity of habitat fragment shape and size increases the diversity of generalist species than isolation.

We therefore conclude that urban planning can improve biodiversity in urban landscapes by managing the size, shape and isolation distance of habitat fragments. Such approaches to urban development can create conditions suitable for avian species persistence in urban landscapes. Large, regular shaped and interconnected habitat fragments are also fundamental to the conservation of avian species in urban landscapes. Future urban development strategies should therefore consider habitat conditions necessary for species persistence, by managing the size, shape and isolation distance of undeveloped grassland and forested areas in urban ecosystems. We suggest further studies that aim to assess the variation of avian species diversity in relation to land use, primary productivity, climatic and topographic variables to assess the pattern of the distribution and assess whether or not further improvements for estimating biodiversity impacts of urban development can be achieved.

## Acknowledgements

We are grateful to Anna Zivumbwa for support in the field.

## Author Contributions

Conceived and designed the study: AM, PT, TM, and NC. Collected the data: TM, PT, NC Analyzed the data: TM. Wrote the manuscript: TM with contributions from AM, PT and NC.

## Supplementary information

**SS 1:** Descriptive statistics of the patch metrics by land use/land cover type

| Patch metrics | Forested |  |  |  | Grasslands |  |  |  | Built-up area |  |  |  |
| --- | --- | --- | --- | --- | --- | --- | --- | --- | --- | --- | --- | --- |
| Size Metrics | Min | Max | Mean | SD | Min | Max | Mean | SD | Min | Max | Mean | SD |
| MeffCUT | 0.01 | 1.70 | 0.61 | 0.58 | 0.05 | 0.63 | 0.33 | 0.21 | 0.00 | 0.21 | 0.10 | 0.07 |
| MeffCBC | 0.14 | 15.58 | 4.38 | 5.17 | 0.06 | 4.98 | 1.07 | 1.45 | 0.10 | 0.21 | 0.25 | 0.35 |
| PLAND | 0.23 | 0.91 | 0.66 | 0.18 | 12.83 | 20.04 | 17.92 | 2.65 | 1.57 | 11.63 | 6.31 | 3.79 |
| NUMB | 4.00 | 127.00 | 42.30 | 40.06 | 9.00 | 226.00 | 67.4 | 2.65 | 108.00 | 400.00 | 288.33 | 96.81 |
| MPS | 2.54 | 197.20 | 68.63 | 67.80 | 2.49 | 86.49 | 26.27 | 25.11 | 0.20 | 4.31 | 0.23 | 1.28 |
| MEDPS | 0.38 | 1.76 | 0.83 | 0.39 | 0.15 | 1.20 | 0.38 | 0.32 | 0.13 | 0.45 | 0.30 | 0.09 |
| PSCOV | 171.9 | 835.84 | 446.05 | 197.66 | 278.24 | 785.23 | 552.5 | 160.95 | 110.12 | 770.14 | 369.09 | 242.05 |
| PSSD | 8.5 | 638.14 | 227.39 | 200.91 | 17.03 | 240.64 | 113.8 | 69.26 | 0.22 | 33.19 | 0.37 | 10.25 |
| **Shape Metrics** |  |  |  |  |  |  |  |  |  |  |  |  |
| ED | 0.02 | 0.12 | 0.05 | 0.03 | 0.02 | 0.12 | 0.05 | 0.03 | 0.11 | 0.33 | 0.20 | 0.06 |
| TE | 19.88 | 117.6 | 45.65 | 33.69 | 16.4 | 66.88 | 34.64 | 15.57 | 20.52 | 73.33 | 42.45 | 17.18 |
| MPE | 0.29 | 4.97 | 2.25 | 1.82 | 0.26 | 1.82 | 0.89 | 0.55 | 0.07 | 0.50 | 0.23 | 0.15 |
| MSI | 1.73 | 2.73 | 2.09 | 0.34 | 1.56 | 2.26 | 1.84 | 0.22 | 1.39 | 1.91 | 1.64 | 0.15 |
| MPAR | 0.17 | 0.35 | 0.25 | 0.06 | 0.18 | 0.44 | 0.35 | 0.08 | 0.27 | 0.45 | 0.34 | 0.05 |
| MSPFD | 1.49 | 1.61 | 1.55 | 0.04 | 1.49 | 1.67 | 1.61 | 0.05 | 1.56 | 1.66 | 1.6 | 0.03 |
| AWMSI | 3.44 | 18.68 | 8.82 | 4.74 | 4.17 | 14.39 | 7.17 | 2.91 | 1.52 | 12.54 | 5.02 | 4.31 |
| AWMPFD | 1.45 | 1.57 | 1.51 | 0.04 | 1.42 | 1.59 | 1.50 | 0.05 | 1.53 | 1.65 | 1.58 | 0.04 |
| **Isolation Metrics** |  |  |  |  |  |  |  |  |  |  |  |  |
| PI | 0.67 | 2.38 | 1.22 | 0.51 | 0.69 | 1.38 | 0.90 | 0.21 | 0.76 | 1.32 | 1.08 | 0.17 |
| OMD | 35.8 | 265.32 | 126.33 | 77.44 | 32.79 | 95.48 | 62.7 | 22.88 | 39.59 | 119.35 | 74.22 | 24.1 |

Note that values for PSSD, MEDPS, MPS, MPE and TE were scaled down by a factor of 1,000. Where: Effective Mesh Size CUT (MeffCUT), Effective Mesh Size CBC (MeffCBC), Percent Landscape Area (PLAND), Number of Patches (NUMB), Mean Patch Size (MPS), Median Patch Size (MEDPS), Patch Size Coefficient of Variation (PSCOV), Patch Size Standard Deviation (PSSD), Edge Density (ED), Total Edge (TE), Mean Patch Edge (MPE), Mean Shape Index (MSI), Mean Perimeter Area Ratio (MPAR), Mean Shape Patch Fractal Dimension (MSPFD), Area Weighted Mean Shape Index (AWMSI), Area Weighted Mean Patch Fractal Dimension (AWMPFD), Proximity Index (PI), Observed Mean Distance (OMD).

**SS2:**
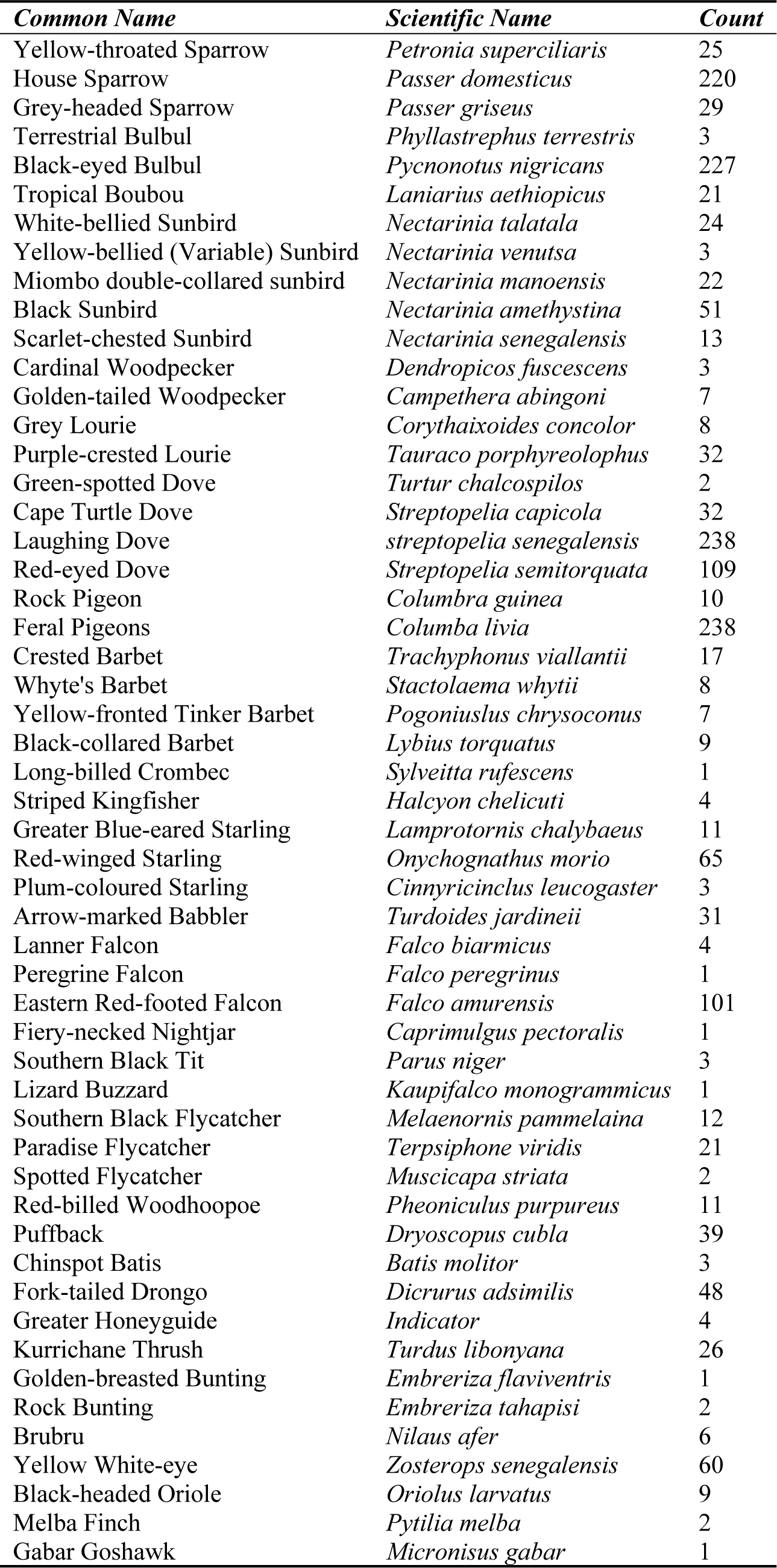
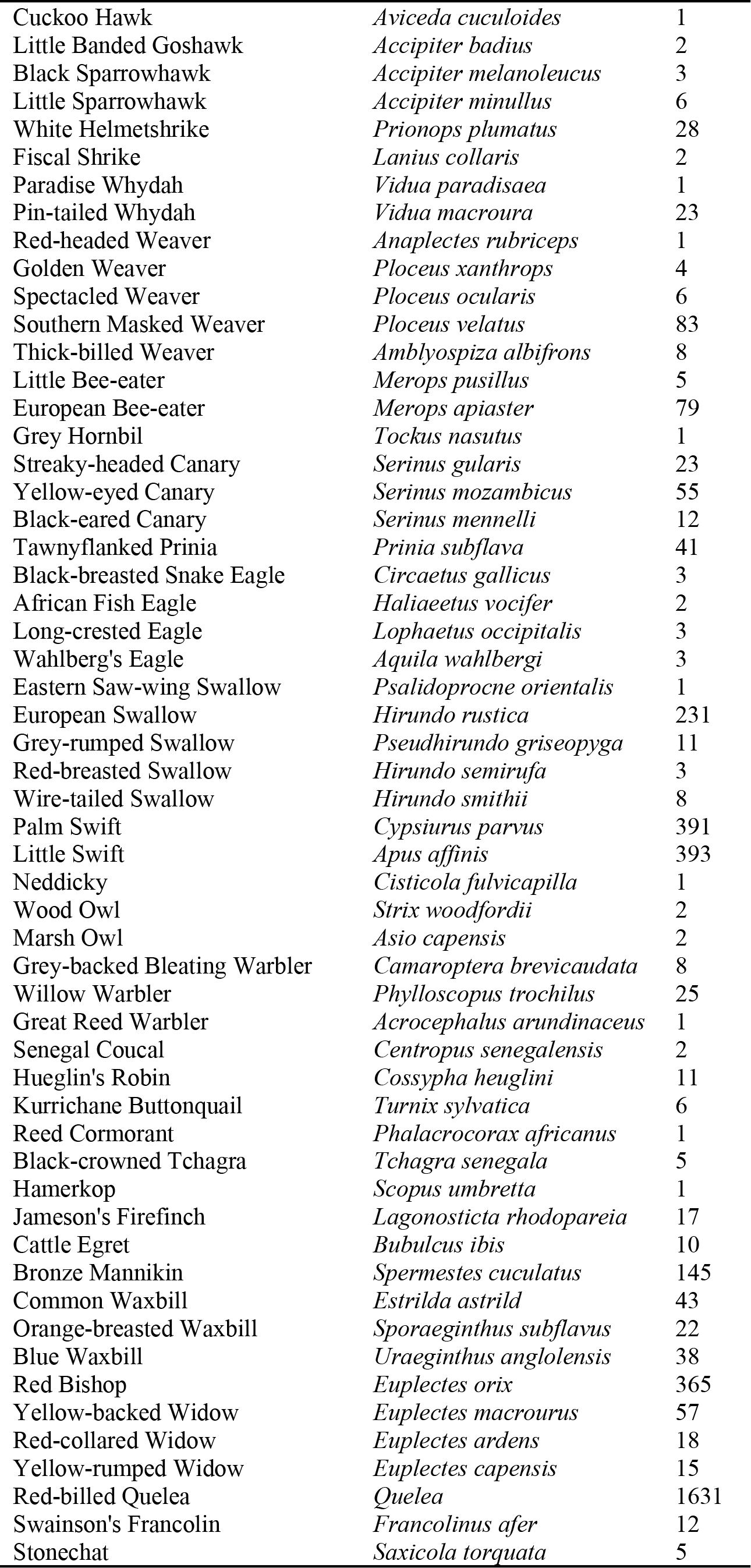
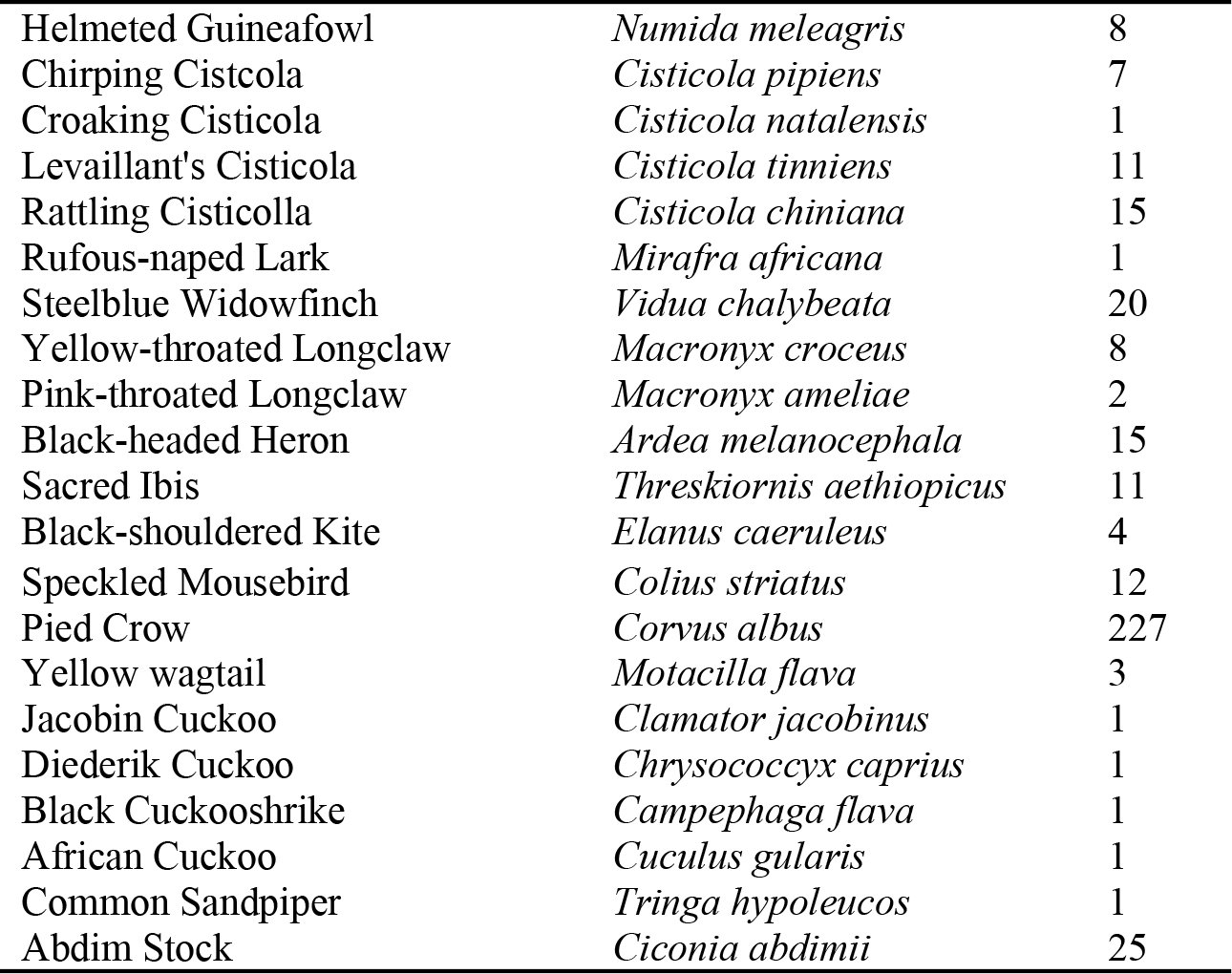
List of avian species recorded in the Harare Metropolitan Region

| *Common Name* | *Scientific Name* | *Count* |
| --- | --- | --- |
| Yellow-throated Sparrow | *Petronia superciliaris* | 25 |
| House Sparrow | *Passer domesticus* | 220 |
| Grey-headed Sparrow | *Passer griseus* | 29 |
| Terrestrial Bulbul | *Phyllastrephus terrestris* | 3 |
| Black-eyed Bulbul | *Pycnonotus nigricans* | 227 |
| Tropical Boubou | *Laniarius aethiopicus* | 21 |
| White-bellied Sunbird | *Nectarinia talatala* | 24 |
| Yellow-bellied (Variable) Sunbird | *Nectarinia venutsa* | 3 |
| Miombo double-collared sunbird | *Nectarinia manoensis* | 22 |
| Black Sunbird | *Nectarinia amethystina* | 51 |
| Scarlet-chested Sunbird | *Nectarinia senegalensis* | 13 |
| Cardinal Woodpecker | *Dendropicos fuscescens* | 3 |
| Golden-tailed Woodpecker | *Campethera abingoni* | 7 |
| Grey Lourie | *Corythaixoides concolor* | 8 |
| Purple-crested Lourie | *Tauraco porphyreolophus* | 32 |
| Green-spotted Dove | *Turtur chalcospilos* | 2 |
| Cape Turtle Dove | *Streptopelia capicola* | 32 |
| Laughing Dove | *streptopelia senegalensis* | 238 |
| Red-eyed Dove | *Streptopelia semitorquata* | 109 |
| Rock Pigeon | *Columbra guinea* | 10 |
| Feral Pigeons | *Columba livia* | 238 |
| Crested Barbet | *Trachyphonus viallantii* | 17 |
| Whyte's Barbet | *Stactolaema whytii* | 8 |
| Yellow-fronted Tinker Barbet | *Pogoniuslus chrysoconus* | 7 |
| Black-collared Barbet | *Lybius torquatus* | 9 |
| Long-billed Crombec | *Sylveitta rufescens* | 1 |
| Striped Kingfisher | *Halcyon chelicuti* | 4 |
| Greater Blue-eared Starling | *Lamprotornis chalybaeus* | 11 |
| Red-winged Starling | *Onychognathus morio* | 65 |
| Plum-coloured Starling | *Cinnyricinclus leucogaster* | 3 |
| Arrow-marked Babbler | *Turdoides jardineii* | 31 |
| Lanner Falcon | *Falco biarmicus* | 4 |
| Peregrine Falcon | *Falco peregrinus* | 1 |
| Eastern Red-footed Falcon | *Falco amurensis* | 101 |
| Fiery-necked Nightjar | *Caprimulgus pectoralis* | 1 |
| Southern Black Tit | *Parus niger* | 3 |
| Lizard Buzzard | *Kaupifalco monogrammicus* | 1 |
| Southern Black Flycatcher | *Melaenornis pammelaina* | 12 |
| Paradise Flycatcher | *Terpsiphone viridis* | 21 |
| Spotted Flycatcher | *Muscicapa striata* | 2 |
| Red-billed Woodhoopoe | *Pheoniculus purpureus* | 11 |
| Puffback | *Dryoscopus cubla* | 39 |
| Chinspot Batis | *Batis molitor* | 3 |
| Fork-tailed Drongo | *Dicrurus adsimilis* | 48 |
| Greater Honeyguide | *Indicator* | 4 |
| Kurrichane Thrush | *Turdus libonyana* | 26 |
| Golden-breasted Bunting | *Embreriza flaviventris* | 1 |
| Rock Bunting | *Embreriza tahapisi* | 2 |
| Brubru | *Nilaus afer* | 6 |
| Yellow White-eye | *Zosterops senegalensis* | 60 |
| Black-headed Oriole | *Oriolus larvatus* | 9 |
| Melba Finch | *Pytilia melba* | 2 |
| Gabar Goshawk | *Micronisus gabar* | 1 |
| Cuckoo Hawk | *Aviceda cuculoides* | 1 |
| Little Banded Goshawk | *Accipiter badius* | 2 |
| Black Sparrowhawk | *Accipiter melanoleucus* | 3 |
| Little Sparrowhawk | *Accipiter minullus* | 6 |
| White Helmetshrike | *Prionops plumatus* | 28 |
| Fiscal Shrike | *Lanius collaris* | 2 |
| Paradise Whydah | *Vidua paradisaea* | 1 |
| Pin-tailed Whydah | *Vidua macroura* | 23 |
| Red-headed Weaver | *Anaplectes rubriceps* | 1 |
| Golden Weaver | *Ploceus xanthrops* | 4 |
| Spectacled Weaver | *Ploceus ocularis* | 6 |
| Southern Masked Weaver | *Ploceus velatus* | 83 |
| Thick-billed Weaver | *Amblyospiza albifrons* | 8 |
| Little Bee-eater | *Merops pusillus* | 5 |
| European Bee-eater | *Merops apiaster* | 79 |
| Grey Hornbil | *Tockus nasutus* | 1 |
| Streaky-headed Canary | *Serinus gularis* | 23 |
| Yellow-eyed Canary | *Serinus mozambicus* | 55 |
| Black-eared Canary | *Serinus mennelli* | 12 |
| Tawnyflanked Prinia | *Prinia subflava* | 41 |
| Black-breasted Snake Eagle | *Circaetus gallicus* | 3 |
| African Fish Eagle | *Haliaeetus vocifer* | 2 |
| Long-crested Eagle | *Lophaetus occipitalis* | 3 |
| Wahlberg's Eagle | *Aquila wahlbergi* | 3 |
| Eastern Saw-wing Swallow | *Psalidoprocne orientalis* | 1 |
| European Swallow | *Hirundo rustica* | 231 |
| Grey-rumped Swallow | *Pseudhirundo griseopyga* | 11 |
| Red-breasted Swallow | *Hirundo semirufa* | 3 |
| Wire-tailed Swallow | *Hirundo smithii* | 8 |
| Palm Swift | *Cypsiurus parvus* | 391 |
| Little Swift | *Apus affinis* | 393 |
| Neddicky | *Cisticola fulvicapilla* | 1 |
| Wood Owl | *Strix woodfordii* | 2 |
| Marsh Owl | *Asio capensis* | 2 |
| Grey-backed Bleating Warbler | *Camaroptera brevicaudata* | 8 |
| Willow Warbler | *Phylloscopus trochilus* | 25 |
| Great Reed Warbler | *Acrocephalus arundinaceus* | 1 |
| Senegal Coucal | *Centropus senegalensis* | 2 |
| Hueglin's Robin | *Cossypha heuglini* | 11 |
| Kurrichane Buttonquail | *Turnix sylvatica* | 6 |
| Reed Cormorant | *Phalacrocorax africanus* | 1 |
| Black-crowned Tchagra | *Tchagra senegala* | 5 |
| Hamerkop | *Scopus umbretta* | 1 |
| Jameson's Firefinch | *Lagonosticta rhodopareia* | 17 |
| Cattle Egret | *Bubulcus ibis* | 10 |
| Bronze Mannikin | *Spermestes cuculatus* | 145 |
| Common Waxbill | *Estrilda astrild* | 43 |
| Orange-breasted Waxbill | *Sporaeeginthus subflavus* | 22 |
| Blue Waxbill | *Uraeginthus angolensis* | 38 |
| Red Bishop | *Euplectes orix* | 365 |
| Yellow-backed Widow | *Euplectes macrourus* | 57 |
| Red-collared Widow | *Euplectes ardens* | 18 |
| Yellow-rumped Widow | *Euplectes capensis* | 15 |
| Red-billed Quelea | *Quelea* | 1631 |
| Swainson's Francolin | *Francolinus afer* | 12 |
| Stonechat | *Saxicola torquata* | 5 |
| Helmeted Guineafowl | *Numida meleagris* | 8 |
| Chirping Cisticola | *Cisticola pipiens* | 7 |
| Croaking Cisticola | *Cisticola natalensis* | 1 |
| Levaillant's Cisticola | *Cisticola tinniens* | 11 |
| Rattling Cisticola | *Cisticola chiniana* | 15 |
| Rufous-naped Lark | *Mirafra africana* | 1 |
| Steelblue Widowfinch | *Vidua chalybeata* | 20 |
| Yellow-throated Longclaw | *Macronyx croceus* | 8 |
| Pink-throated Longclaw | *Macronyx ameliae* | 2 |
| Black-headed Heron | *Ardea melanocephala* | 15 |
| Sacred Ibis | *Threskiornis aethiopicus* | 11 |
| Black-shouldered Kite | *Elanus caeruleus* | 4 |
| Speckled Mousebird | *Colius striatus* | 12 |
| Pied Crow | *Corvus albus* | 227 |
| Yellow wagtail | *Motacilla flava* | 3 |
| Jacobin Cuckoo | *Clamator jacobinus* | 1 |
| Diederik Cuckoo | *Chrysococcyx caprius* | 1 |
| Black Cuckooshrike | *Campephaga flava* | 1 |
| African Cuckoo | *Cuculus gularis* | 1 |
| Common Sandpiper | *Tringa hypoleucos* | 1 |
| Abdim Stock | *Ciconia abdimii* | 25 |

## Possible Reviewers

1. Seth Mago, Director, Urban Wildlife Institute, Lincoln Park Zoo,
2. Wendy McWilliam, Senior Lecturer, Lincoln University, NZ,
3. Amanda D Rodwald, Professor; Director of Conservation Science, Cornell Lab of Ornithology

